# A dynamic bioenergetic model for coral-*Symbiodinium* symbioses and coral bleaching as an alternate stable state

**DOI:** 10.1101/120733

**Authors:** Ross Cunning, Erik B. Muller, Ruth D. Gates, Roger M. Nisbet

## Abstract

Coral reef ecosystems owe their ecological success–and vulnerability to climate change–to the symbiotic metabolism of corals and *Symbiodinium* spp. The urgency to understand and predict the stability and breakdown of these symbioses (i.e., coral ‘bleaching’) demands the development and application of theoretical tools. Here, we develop a dynamic bioenergetic model of coral-*Symbiodinium* symbioses that demonstrates realistic steady-state patterns in coral growth and symbiont abundance across gradients of light, nutrients, and feeding. Furthermore, by including a mechanistic treatment of photo-oxidative stress, the model displays dynamics of bleaching and recovery that can be explained as transitions between alternate stable states. These dynamics reveal that ‘healthy’ and ‘bleached’ states correspond broadly to nitrogen- and carbon-limitation in the system, with transitions between them occurring as integrated responses to multiple environmental factors. Indeed, a suite of complex emergent behaviors reproduced by the model (e.g., bleaching is exacerbated by nutrients and attenuated by feeding) suggests it captures many important attributes of the system; meanwhile, its modular framework and open source R code are designed to facilitate further problem-specific development. We see significant potential for this modeling framework to generate testable hypotheses and predict integrated, mechanistic responses of corals to environmental change, with important implications for understanding the performance and maintenance of symbiotic systems.

## Introduction

The nutritional exchange between corals and *Symbiodinium* directly underlies the capacity of corals to build coral reef ecosystems, worth trillions of US Dollars annually (Costanza, Groot, and Sutton 2014). However, the complex symbiotic metabolism of corals is vulnerable to disruption by numerous anthropogenic environmental perturbations, jeopardizing their future persistence. In order to understand and predict responses of corals to complex changes in their environment, a mechanistic understanding of how multiple interacting factors drive the individual and emergent physiology of both symbiotic partners is necessary. Such a task is well suited for theoretical modeling frameworks such as Dynamic Energy Budget (DEB) theory (Kooijman 2010), although the complexity of such theory makes these efforts inaccessible to many biologists (Jager, Martin, and Zimmer 2013). In order to bridge this gap, we present here a simplified dynamic bioenergetic model for coral-*Symbiodinium* symbioses that aims to mechanistically integrate the impacts of complex environmental change on the physiological performance of reef corals, including responses to environmental stress.

In reef coral symbioses, intracellular *Symbiodinium* translocate photosynthetically fixed carbon to support coral metabolism, while the animal host provides access to inorganic nutrients and carbon dioxide (Muscatine and Porter 1977). Previous application of DEB theory to this syntrophic system (Muller et al. 2009) demonstrated a stable symbiotic relationship and qualitatively realistic growth and biomass ratios across gradients of ambient irradiance, nutrients, and food. This model (as well as the present work) assumes that 1) *Symbiodinium* has priority access to fixed carbon through photosynthesis, 2) the coral animal has priority access to nitrogen through contact with seawater and feeding on prey, and 3) each partner shares with the other only what it cannot use for its own growth. In its simplest form, this principle of sharing the surplus is sufficient to describe the dynamics of diverse syntrophic organs and organisms (e.g., trees, duckweeds, corals), suggesting the mechanism is mathematically and evolutionarily robust (Nisbet et al., in prep.).

While the formal DEB model of Muller et al. (2009) represents the most significant theoretical contribution in coral symbiosis research to date, we aim to strengthen the role of theory and broaden its potential application in three primary ways:

1. *Develop a new module of photo-oxidative stress*. Of primary interest to coral biologists and ecologists is symbiosis dysfunction under environmental stress, resulting in coral “bleaching”–the loss of algal symbionts from the association (Jokiel and Coles 1977). Bleaching is thought to be triggered by photo-oxidative stress in *Symbiodinium* (Weis 2008), which has been modeled previously as a response to absolute external irradiance (Eynaud, Nisbet, and Muller 2011). However, this response may also depend on self-shading by other symbionts (Enríquez, Méndez, and Iglesias-Prieto 2005), CO_2_ availability at the site of photosynthesis (Wooldridge 2009), and other non-photochemical quenching mechanisms (Roth 2014). We incorporate these features into a new photo-oxidative stress module linking overreduction of the photosynthetic light reactions to downstream impacts of photoinhibition and photodamage. Importantly, this formulation introduces a link between CO_2_-limitation of photosynthesis and bleaching, and potential synergistic roles for heterotrophy and nutrient availability in influencing bleaching responses (Wooldridge 2014b)
2. *Reduce theoretical and mathematical complexity*. Following the logic of Jager, Martin, and Zimmer (2013), we exclude certain features of formal DEB theory in order to capture behaviors of interest with the simplest possible formulation. Here, we present a model without reserves, maturity, or reproduction (see Kooijman 2010). This formulation precludes modeling the full life cycle of corals as reproduction, larval stages, and metamorphosis are not considered, but greatly reduces theoretical complexity and parameter numbers, which is advantageous given the relative paucity of data for corals. Moreover, dynamics of the symbiosis (i.e., changes in symbiont to host biomass, including bleaching and recovery) and coral biomass growth are efficiently captured with this simpler formulation, which also increases accessibility for biologists and ecologists without requiring expertise in DEB theory.
3. *Provide well-documented, open source code*. In order to facilitate the continued development and application of theoretical modeling tools for coral symbioses, we provide access to the model in the form of an R package called coRal github.com/jrcunning/coRal, which users may install to run and visualize model simulations. With an accessible and modular framework, we envision this code base as a resource for further development by the scientific community to include additional complexity and problem-specific components. We chose R (R Core Team 2014), an open source programming language in common use by biologists and ecologists, to reach the widest possible audience with this work.

With these as our primary motivations, we describe a simplified approach to bioenergetic modeling of coral-*Symbiodinium* symbioses that dynamically integrates the influences of external irradiance, nutrients, and prey availability on coral growth and symbiosis dynamics (i.e., changes in symbiont:host biomass ratios), allowing for the possibility of coral bleaching in response to photooxidative stress. An emergent finding of this work is that coral bleaching can be interpreted as an alternate stable state of the symbiotic system, which provides a new framework for understanding the mechanisms that drive a coral into a bleached state, as well as those that facilitate recovery. In the following sections, we describe and provide rationale for the model structure, demonstrate a range of steady state and dynamic behaviors that are consistent with observed phenomena, and discuss new insights from this work in understanding responses of coral*-Symbiodinium* symbioses to environmental change.

## Model description

In this dynamical system, both the coral host and algal symbiont acquire and use carbon and nitrogen to construct biomass. The symbiont fixes carbon through photosynthesis and receives nitrogen shared by the host, while the host acquires nitrogen from the environment and receives carbon shared by the symbiont. A graphical representation of the model is presented in Fig. 1, and each model flux and parameter is defined in Tables 1 and 2, respectively. We use C-moles (C-mol) as the unit of biomass for consistency with the rigorous mass balance of DEB theory: 1 C-mol is equivalent to the amount of biomass containing 1 mole of carbon atoms. Host biomass (expressed as C-mol H), symbiont biomass (expressed as C-mol S), and prey biomass (expressed as C-mol X) have fixed, but different, molar N:C ratios (Table 2). Biomass is produced from carbon and nitrogen by synthesizing units (SU), which are mathematical specifications of the formation of a product from two substrates; we use the “parallel complementary” formulation of Kooijman (2010) (page 105, Fig. 3.7) to specify these fluxes. The two state variables of this system are symbiont biomass and coral biomass; because resources are acquired proportionally to surface area, and surface area is proportional to volume, biomass increases exponentially during growth (indeed, corals grow exponentially (Bak 1976)).

**Figure 1:**
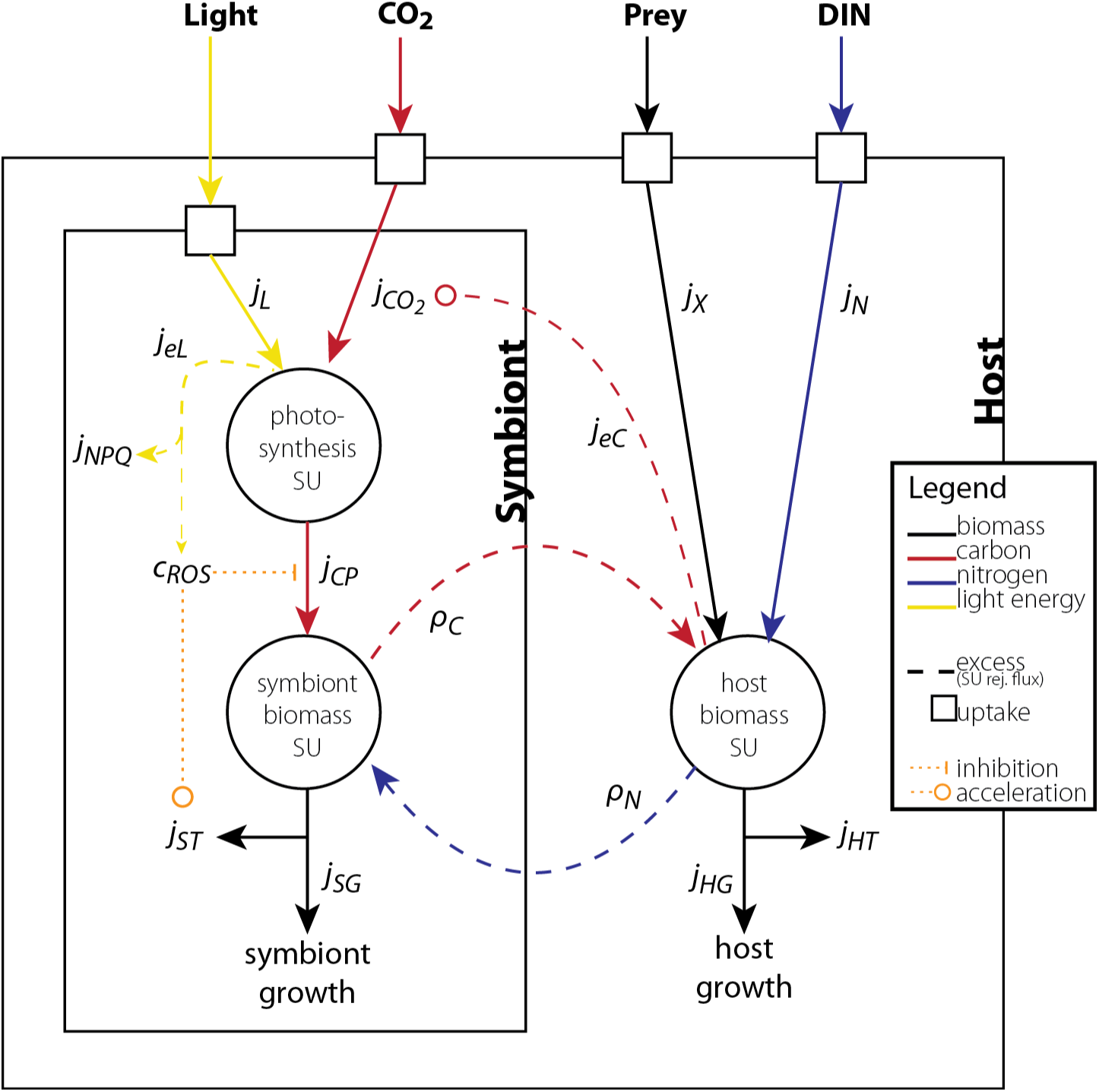
Graphical representation of coral-algal symbiosis model. Light, CO_2_, prey, and DIN are acquired from the external environment proportional to the biomass of the partner indicated by the black box for uptake. Mass fluxes (see Table 1 for definitions) are represented by *j*’s with subscripts indicating the type of mass, and in some cases the process (e.g., *j*_*CP*_ is the flux of carbon produced by photosynthesis), and *ρ*’s indicate fluxes that are shared by one partner with the other. Parallel complementary synthesizing units (SUs) are represented by large circles, and rejection fluxes from these SUs are indicated by dashed lines. *c*_*ROS*_ is a proportional rate that impacts other model fluxes by inhibition or acceleration; likewise, *j*_*eC*_ accelerates the rate of *j*_*CO*2_. Recycling fluxes are not shown for clarity (but see Table 1 for definitions).

**Table 1.** Model fluxes (mass-specific). Units are explained in the text.

| Symbol | Description | Units | Eq. no. |
| --- | --- | --- | --- |
| $j_X$ | Prey assimilation (feeding) rate | C-mol X C-mol H<sup>-1</sup> d<sup>-1</sup> | 3 |
| $j_N$ | Nitrogen uptake rate | mol N C-mol H<sup>-1</sup> d<sup>-1</sup> | 4 |
| $j_{HG}$ | Host biomass formation rate | C-mol H C-mol H<sup>-1</sup> d<sup>-1</sup> | 5 |
| $j_{HT}$ | Host biomass turnover rate | C-mol H C-mol H<sup>-1</sup> d<sup>-1</sup> | 6 |
| $r_{NH}$ | Recycled nitrogen from host turnover | mol N C-mol H<sup>-1</sup> d<sup>-1</sup> | 7 |
| $\rho_N$ | Nitrogen shared with the symbiont | mol N C-mol H<sup>-1</sup> d<sup>-1</sup> | 8 |
| $j_{eC}$ | Excess carbon used to activate host CCMs | mol C C-mol H<sup>-1</sup> d<sup>-1</sup> | 9 |
| $j_{CO_2}$ | CO<sub>2</sub> input to photosynthesis | mol CO<sub>2</sub> C-mol H<sup>-1</sup> d<sup>-1</sup> | 10 |
| $j_L$ | Light absorption rate | mol photons C-mol S<sup>-1</sup> d<sup>-1</sup> | 12 |
| $r_{CH}$ | Recycled CO<sub>2</sub> from host | mol CO<sub>2</sub> C-mol H<sup>-1</sup> d<sup>-1</sup> | 13 |
| $r_{CS}$ | Recycled CO<sub>2</sub> from symbiont | mol CO<sub>2</sub> C-mol S<sup>-1</sup> d<sup>-1</sup> | 14 |
| $j_{CP}$ | Photosynthesis rate | mol C C-mol S<sup>-1</sup> d<sup>-1</sup> | 15 |
| $j_{eL}$ | Light energy in excess of photochemistry | mol photons C-mol S<sup>-1</sup> d<sup>-1</sup> | 16 |
| $j_{NPQ}$ | Total capacity of NPQ | mol photons C-mol S<sup>-1</sup> d<sup>-1</sup> | 17 |
| $c_{ROS}$ | ROS production proportional to baseline | – | 18 |
| $r_{NS}$ | Recycled nitrogen from symbiont turnover | mol N C-mol S<sup>-1</sup> d<sup>-1</sup> | 19 |
| $j_{SG}$ | Symbiont biomass formation rate | C-mol S C-mol S<sup>-1</sup> d<sup>-1</sup> | 20 |
| $\rho_C$ | Fixed carbon shared with host | mol C C-mol S<sup>-1</sup> d<sup>-1</sup> | 21 |
| $j_{ST}$ | Symbiont biomass turnover rate | C-mol S C-mol S<sup>-1</sup> d<sup>-1</sup> | 22 |

**Table 2.** Model parameters (units explained in the text). Justification and/or derivation of each parameter value along with supporting references are provided in the Supplementary Information.

| Symbol | Description | Units | Value |
| --- | --- | --- | --- |
| $n_{NH}$ | N:C molar ratio in host biomass | – | 0.18 |
| $n_{NS}$ | N:C molar ratio in symbiont biomass | – | 0.13 |
| $n_{NX}$ | N:C molar ratio in prey biomass | – | 0.2 |
| $j_{HT}^0$ | Maintenance rate of host biomass | C-mol H C-mol H<sup>-1</sup> d<sup>-1</sup> | 0.03 |
| $j_{ST}^0$ | Maintenance rate of symbiont biomass | C-mol S C-mol S<sup>-1</sup> d<sup>-1</sup> | 0.03 |
| $\sigma_{NH}$ | Proportion N turnover recycled in host | – | 0.9 |
| $\sigma_{CH}$ | Proportion host metabolic CO<sub>2</sub> recycled to photosynthesis | – | 0.1 |
| $\sigma_{NS}$ | Proportion N turnover recycled in symbiont | – | 0.9 |
| $\sigma_{CS}$ | Proportion symbiont metabolic CO<sub>2</sub> recycled to photosynthesis | – | 0.9 |
| $j_{Xm}$ | Maximum prey assimilation rate from host feeding | C-mol X C-mol H<sup>-1</sup> d<sup>-1</sup> | 0.13 |
| $K_X$ | Half-saturation constant for prey assimilation | C-mol X L<sup>-1</sup> | 10<sup>-6</sup> |
| $j_{Nm}$ | Maximum host DIN uptake rate | mol N C-mol H<sup>-1</sup> d<sup>-1</sup> | 0.035 |
| $K_N$ | Half-saturation constant for host DIN uptake | mol N L<sup>-1</sup> | 1.5 × 10<sup>-6</sup> |
| $k_{CO_2}$ | Efficacy of CO<sub>2</sub> delivery to photosynthesis by host CCMs | mol CO<sub>2</sub> mol C<sup>-1</sup> | 10 |
| $j_{HGm}$ | Maximum specific growth rate of host | C-mol H C-mol H<sup>-1</sup> d<sup>-1</sup> | 1 |
| $y_{CL}$ | Quantum yield of photosynthesis | mol C mol photons<sup>-1</sup> | 0.1 |
| $y_C$ | Yield of biomass formation from carbon | C-mol mol C<sup>-1</sup> | 0.8 |
| $\bar{a}^*$ | Effective light-absorbing cross-section of symbiont | m<sup>2</sup> C-mol S<sup>-1</sup> | 1.34 |
| $k_{NPQ}$ | NPQ capacity of symbiont | mol photons C-mol S<sup>-1</sup> d<sup>-1</sup> | 112 |
| $k_{ROS}$ | Excess photon energy that doubles ROS production, relative to baseline levels | mol photons C-mol S<sup>-1</sup> d<sup>-1</sup> | 80 |
| $j_{CPm}$ | Maximum specific photosynthesis rate of symbiont | mol C C-mol S<sup>-1</sup> d<sup>-1</sup> | 2.8 |
| $j_{SGm}$ | Maximum specific growth rate of symbiont | C-mol S C-mol S<sup>-1</sup> d<sup>-1</sup> | 0.25 |
| $b$ | Scaling parameter for bleaching response | – | 5 |

Environmental stress is implemented in the form of photooxidative stress, which is thought to be a primary trigger of coral bleaching (Lesser 1997; Weis 2008; Wooldridge 2009). To simulate bleaching, we model the absorption and quenching of light energy by photochemistry and non-photochemical quenching, and the responses that occur (i.e., photoinhibition, photodamage, and symbiont loss) when these quenching capacities are overwhelmed. While bleaching in response to high light alone has been observed experimentally (Schutter et al. 2011; Downs et al. 2013), mass coral bleaching events occur concurrently with high temperature (Hoegh-Guldberg 1999). Thus, it is important to justify our consideration of light as the primary stressor. In reality, light and temperature interact synergistically (Coles and Jokiel 1978; Jones et al. 1998), and in fact, any stressor that disrupts the quenching of light energy may lead to bleaching (Wooldridge 2010; Baker and Cunning 2015). This is because the proximate cause of photo-oxidative stress is excess excitation energy, but the upstream events that lead to this situation may be diverse. Indeed, elevated temperature may inhibit Rubisco functioning (Jones et al. 1998) and the repair of the D1 protein in photosystem II (Warner, Fitt, and Schmidt 1999), which reduces the capacity of photochemical quenching and leads to an excess of light energy. In this way, elevated temperature serves to reduce the threshold above which light stresses the system (Hoegh-Guldberg 1999); importantly, light is still the proximate stressor. Therefore, we omitted temperature from the model to maintain a desired level of simplicity, while still allowing photooxidative stress and bleaching to be simulated with biological realism in response to light.

### State equations

The balance equations for symbiont (*S*) and host (*H*) biomass are expressed as “specific” rates, i.e. rates per unit of symbiont and host biomass, respectively:

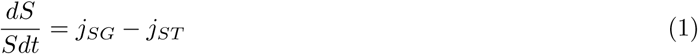

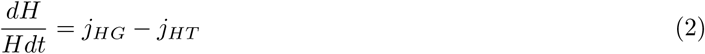

The specific biomass growth and turnover rates that define these balance equations are produced by combinations of the individual model fluxes (see Table 1 for definitions and units), which are each expressed as mass-specific rates (e.g., per C-mole of symbiont or host biomass per day). When necessary, conversions between symbiontmass-specific and host-mass-specific rates are accomplished by multiplying or dividing by the symbiont:host biomass ratio.

### Coral animal fluxes

The coral animal acquires both carbon and nitrogen from feeding on prey from the environment. Assimilation from feeding is specified by Michaelis-Menten kinetics (i.e., a Holling type II function) with a maximum rate of *j*_*Xm*_ and half-saturation constant *K*_*X*_:

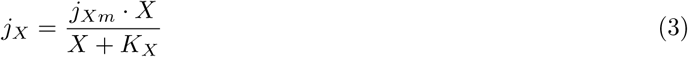

Additionally, the coral animal acquires dissolved inorganic nitrogen (DIN) from the surrounding seawater, which is assumed to represent ammonium, the primary form utilized by corals (Yellowlees, Rees, and Leggat 2008). This gives the host (rather than the symbiont) priority in nitrogen utilization; this capacity is supported by experimental evidence (Wang and Douglas 1998) and is consistent with the spatial arrangment of the partners, where the host is in direct contact with the external environment. The uptake of nitrogen from the environment is thus specified by Michaelis-Menten kinetics using a maximum uptake rate *j*_*Nm*_ and half-saturation constant *K*_*N*_:

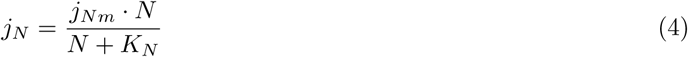

Coral biomass formation is then specified by a parallel complementary SU (formula in Fig. 3.7 of Kooijman (2010)]) that combines carbon and nitrogen, according to:

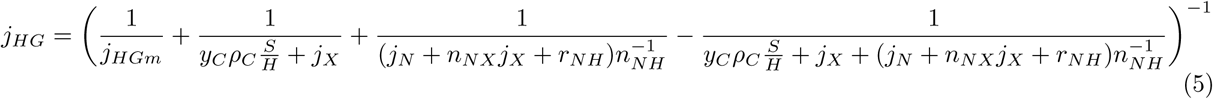
where *ρ*_*C*_ is fixed carbon shared by the symbiont (see Eq. 21), and *r*_*NH*_ is recycled nitrogen liberated by host biomass turnover (see Eq. 7). The parameter *y*_*C*_ specifies the yield of biomass from organic carbon, which we take to be 0.8 to satisfy redox balance (see Muller et al. (2009)).

Host biomass turnover is equal to the specific maintenance rate of host biomass,

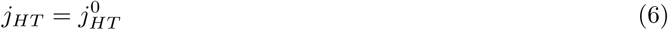
and the specific flux of nitrogen that is recycled to the host biomass SU is calcluated as:

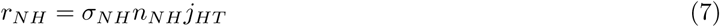

The amount of nitrogen input to the coral biomass SU in excess of what is actually consumed in biomass formation (i.e., surplus nitrogen, or the rejection flux^1^ of the SU) is then made available to the symbiont:

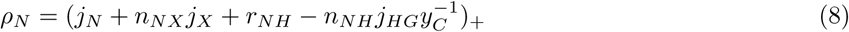

Due to the inherent inefficiency of the parallel complementary SU formulation, there is always some nitrogen shared with the symbiont even when coral biomass formation is strongly nitrogen-limited. Likewise, there is always a non-zero rejection flux of excess carbon from the coral biomass SU. The carbon rejected from this SU reflects the amount of excess fixed carbon available to the host that is not used in biomass formation:

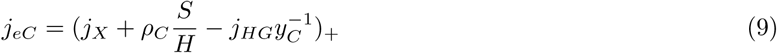

This flux, *j*_*eC*_, is assumed to be available to the host as a respiratory substrate to support energetically-demanding processes; of particular importance is the host’s active carbon concentrating mechanisms (CCMs) that supply CO_2_ for symbiont photosynthesis (Wooldridge 2013; Hopkinson, Tansik, and Fitt 2015). We therefore specify *j*_*CO*_2__ as the host-mediated delivery of CO_2_ to photosynthesis that encompasses potentially diverse CCMs, including active transport of bicarbonate, carbonic anhydrase-catalyzed conversion of bicarbonate to CO_2_ to promote diffusion toward the symbiont (Tansik, Fitt, and Hopkinson 2015), and acidification of the symbiosome to increase localized CO_2_ concentrations around the symbiont (Barott et al. 2014). Since these active CCMs require energetic input by the host, we define *j*_*CO*_2__ as proportional to *j*_*eC*_, assuming that some of this carbon is respired to energize the CCMs. This formulation means that the symbiont indirectly ensures its own CO_2_ supply by providing fixed carbon (=energy) to the host (Wooldridge 2013). The parameter *k*_*CO*_2__ scales the efficacy of host CCMs, which enables the comparison of different rates of CO_2_ delivery that may characterize different coral species (Wooldridge 2014a). The active input of CO_2_ to the photosynthesis SU is therefore specified as:

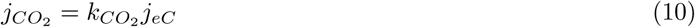

### Symbiodinium *fluxes*

The symbiont produces fixed carbon through photosynthesis, a process represented here by a single SU with two substrates: light (photons) and inorganic carbon (CO_2_). The amount of light absorbed by the symbiont depends on the scalar irradiance at the site of light absorption, which is modified substantially relative to external downwelling irradiance owing to multiple scattering by the coral skeleton and self-shading by surrounding symbionts (Enríquez, Méndez, and Iglesias-Prieto 2005; Marcelino et al. 2013). We used data from Marcelino et al. (2013) to empirically derive an amplification factor, A, indicating the ratio of internal scalar irradiance to external downwelling irradiance as a function of symbiont density (S:H biomass), which is specified as:

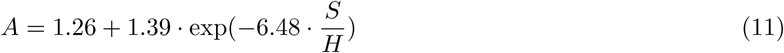

This amplification factor is then multiplied by the external downwelling irradiance *L* and a parameter representing the effective light-absorbing surface area of symbiont biomass *ā*\* to specify the total light absorption:

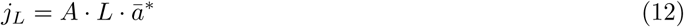

CO_2_ arrives at the photosynthesis SU from multiple sources: in addition to the CO_2_ actively supplied by the host through its CCMs (*j*_*CO*_2__; Eq. 10), we assume a fixed proportion *σ*_*CH*_ of metabolic CO_2_ generated by the host from both biomass turnover and formation is passively available to the photosynthesis SU, according to:

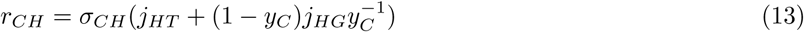
along with a fixed proportion of CO_2_ generated by symbiont biomass turnover^2^ and formation:

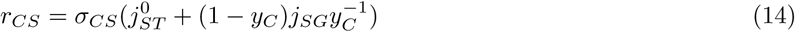

Fixed carbon is then produced by the photosynthesis SU according to:

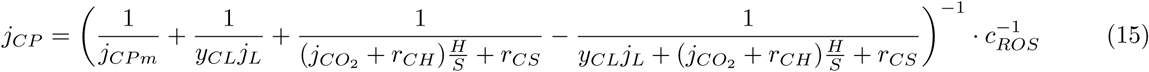
where *j*_*CPm*_ is the maximum specific rate of photosynthesis, and *c*_*ROS*_ is the relative rate of reactive oxygen species production (see Eq. 18). Dividing the photosynthetic rate by *c*_*ROS*_ causes a decline in response to photooxidative stress at high light levels, and the emergent outcome of this SU formulation demonstrates a classic photoinhibition response (Fig. S2).

Light energy absorbed in excess of what is used to fix carbon is specified by the SU rejection flux, according to:

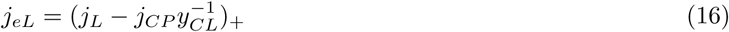

This excess light energy must be quenched by alternative pathways in order to prevent photooxidative damage (Powles 1984). *Symbiodinium* utilize a variety of pathways for non-photochemical quenching (NPQ; Roth 2014), which we collect in a total NPQ capacity specified as a parameter of the symbiont (*k*_*NPQ*_). The NPQ flux *j*_*NPQ*_ is then specified as a single-substrate SU formula with a maximum of *k*_*NPQ*_:

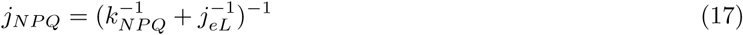

If light energy further exceeds the capacity of both photochemistry and NPQ, then reactive oxygen species (ROS) are produced. We represent this as a relative quantity *c*_*ROS*_, which takes a value of 1 when all light energy is quenched by photochemistry and NPQ, and increases as the amount of excess excitation energy increases, specified as:

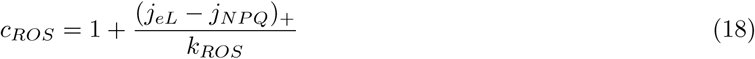
where *k*_*ROS*_ is a parameter of the symbiont that determines the rate of ROS production (specifically, the amount of excess excitation energy that doubles ROS production relative to baseline levels). Importantly, *c*_*ROS*_ is specified here not as a function of absolute external light, but rather the amount of excess light energy after accounting for quenching by carbon fixation and NPQ. A direct consequence of this formulation is that CO_2_-limitation of photosynthesis can lead to ROS production, an important mechanism (Butow et al. 1998; Wooldridge 2009) that was not captured by previous representations of photooxidative stress (Eynaud, Nisbet, and Muller 2011). With this single SU, both the light and dark reactions of photosynthesis are represented, allowing for sink-limitation (i.e., CO_2_-limitation) to cause overreduction of the electron transport chain and ROS production.

Carbon fixed by photosynthesis (*j*_*CP*_; Eq. 15) is then combined with nitrogen shared by the host (*ρ*_*N*_; Eq. 8) and nitrogen recycled from symbiont biomass turnover^3^

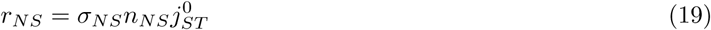
to build new symbiont biomass, following the SU equation:

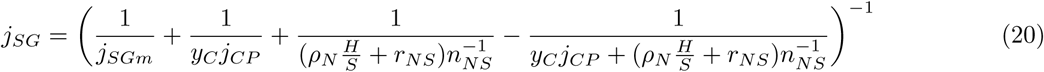

The rejection flux of carbon from this SU represents the amount of fixed carbon produced by photosynthesis in excess of what can be used to produce symbiont biomass; this surplus, *ρ*_*C*_, is translocated to the coral host:

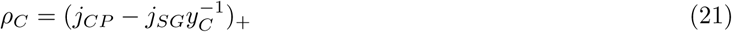

Nitrogen rejected by the symbiont biomass SU, which has already been rejected by the host biomass SU, cannot be used by either partner and is thus lost to the environment.

Symbiont biomass turnover includes a component of constant turnover specified by the parameter 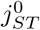, representing fixed maintenance costs, plus a component that scales with the magnitude of ROS production.

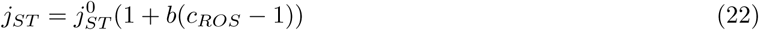

This second component of symbiont biomass loss represents both photodamage and/or symbiont expulsion (i.e., bleaching), both of which occur in response to high levels of ROS production. The parameter *b* is included to scale biomass loss due to bleaching in response to ROS.

To aid in visualizing model results, we calculated values to indicate the degree to which product formation at an SU was limited by availability of either of its two substrates using the formula

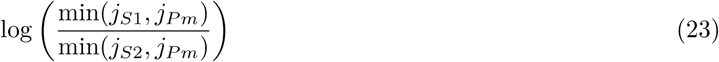
where *j*_*S*1_ and *j*_*S*2_ are the specific input fluxes of the two substrates and *j*_*Pm*_ is the maximum specific product formation rate, in units of Cmol Cmol^-1^ d^-1^. When both substrate input fluxes are higher than what can be used at the maximum production rate, this limitation coefficient is zero, implying that neither substrate is limiting production.

### Numerical analysis

The model dynamics are specified by the differential equations (1-2) that impose biomass balance for host and symbiont and by a set of coupled non-linear algebraic equations (3-22) that define fluxes. Several of these fluxes are defined *implicitly*; for example, the rejection fluxes of carbon and nitrogen from the symbiont and host biomass SUs, respectively, act as reciprocal input fluxes to the other SU. Similarly, the photosynthesis SU receives CO_2_ at a rate proportional to the carbon rejection flux from the host biomass SU, and the rejection flux of excitation energy from the photosynthesis SU acts to reduce its own production through photoinhibition. Without further assumptions, however, the dynamical system is not always unambiguously defined because for some combinations of parameters and environmental forcing functions the system of algebraic equations has more than one solution with all fluxes non-negative (see results below). In such circumstances, the right hand side of the differential equations (1) and (2) is not uniquely defined even when *S* and *H* are specified. We resolved this problem by *defining the dynamical system* as the limit as a time step Δ*t* → 0 of a discretized system corresponding to Euler integration of the differential equations, with those fluxes that represent flows of elemental matter implemented by assuming that transfer of material between components of the system takes one time step. Thus, for example, CO_2_ rejected from the host SU at time *t* arrives at the photosynthesis SU at time *t* + Δ*t*.

Simulations using the discretized scheme were performed using R code developed in the coRal R package github.com/jrcunning/coRal. By experimentation, we found that a time step of 0.1 days gave adequate precision for most simulations (including used to generate Figs. 2-8 in this paper). For steady state estimations, simulations were run until the changes in specific growth rate of the host and the S:H biomass ratio were less than 1e-5 per time step. In regions of state space where very slow transient dynamics could be expected (i.e. near bifurcation points), sample steady state calculations were verified using MATHEMATICA code for numerical root finding (function FindRoot) with the code written independently by a coauthor without reference to the R code. All of the R code for the simulations and figures presented in this paper can be found in the accompanying data repository at github.com/jrcunning/coRal-analysis.

## Steady state behavior

In a constant environment, the system ultimately reaches a steady state of exponential growth or decline. However, under some conditions, either of these outcomes may occur depending on initial values of symbiont and host biomass, indicating the presence of alternate stable states (Fig. 2). The mechanism that produces these alternate stable states is the positive feedback between carbon-limitation of the host and CO_2_-limitation of photosynthesis: if symbiont biomass is initially very low (i.e., a “bleached” coral), very little carbon is fixed, and the system cannot escape this positive feedback and cannot grow (unless feeding is sufficiently high). However, if symbiont biomass is initially high (i.e., a “healthy” coral), then the system remains in a nitrogen-limited state with positive growth. For practical purposes, this section of the manuscript considers only positive growth steady states under constant environments; subsequently, we explore how environmental forcing may cause the system to switch between alternate stable states, which we interpret in the context of coral bleaching (see “Coral Bleaching and Recovery”, below).

**Figure 2:**
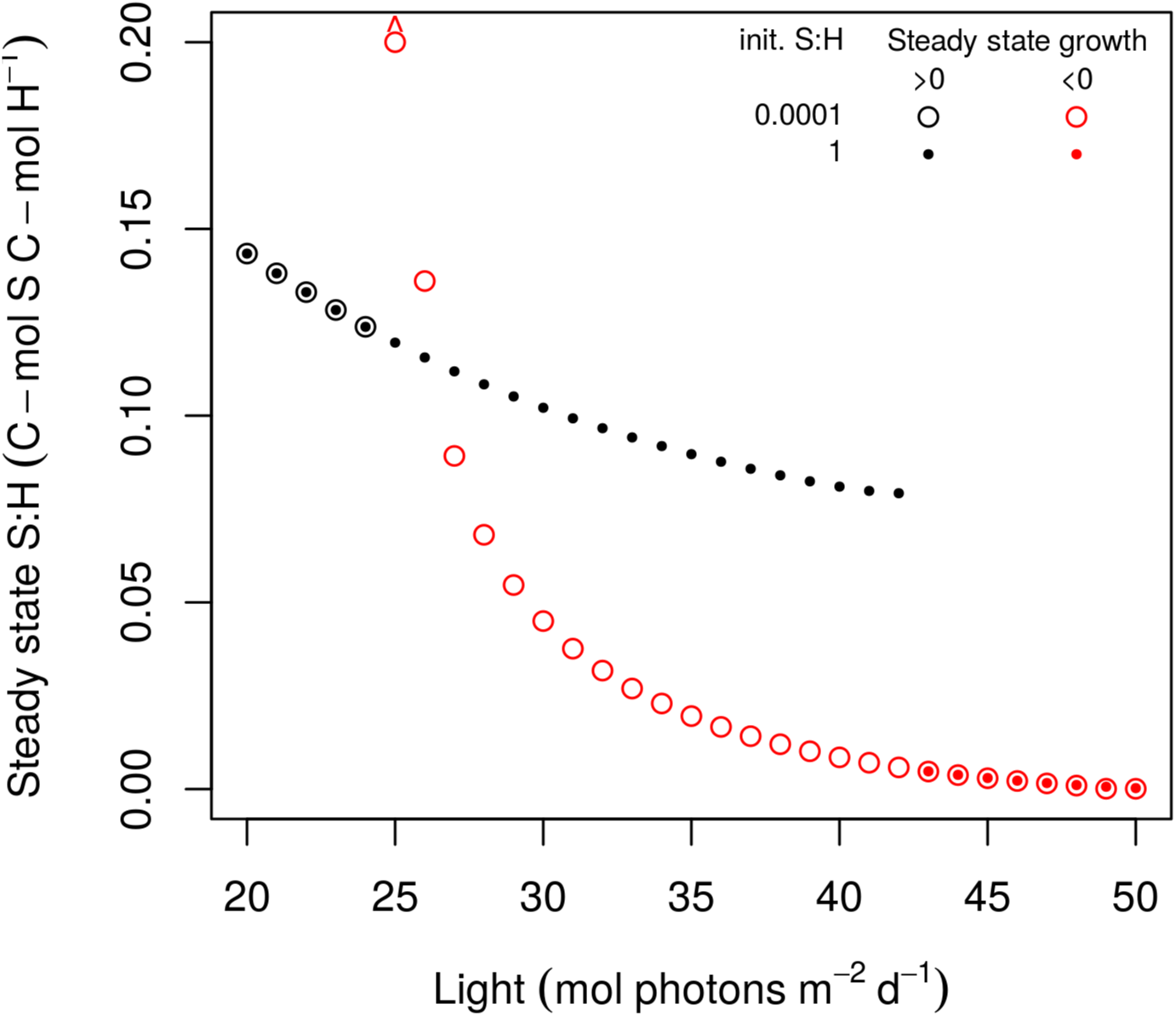
Alternate stable states in S:H biomass and growth across a light gradient. Alternate stable states occur between ∼25-42 mol photons m^-2^ d^-1^ under these conditions (DIN=1e-7 mol N L^-1^; X=1e-7 C-mol X L^-1^), depending on whether initial S:H is high (1, closed circles), representing a healthy coral, or low (0.0001, open circles), representing a bleached coral. Arrow above point at L=25 indicates a S:H ratio beyond the axis range; this ‘overshoot’ phenomenon, in which initially bleached corals may achieve high S:H ratios while remaining in a carbon-limited state is discussed in the *Coral Bleaching and Recovery* section.

To analyze positive-growth steady state behavior, we ran the model to steady state across gradients of external irradiance and nutrients (Fig. 3), which revealed patterns consistent with observed phenomena in corals. Predicted growth rates are low at low light and DIN (∼0.01 d^-1^), and begin increasing as both of these factors increase (Fig. 3A). Low light limits photosynthetic rates, resulting in less fixed carbon shared with the host and an associated increase in the symbiont to host biomass ratio (Fig. 3B). In agreement with this trend are many observations of negative correlation between irradiance and symbiont density (Stimson 1997; Brown et al. 1999; Fitt et al. 2000; Titlyanov et al. 2001). As higher light alleviates light-limitation of photosynthesis, host growth becomes less carbon-limited.

**Figure 3:**
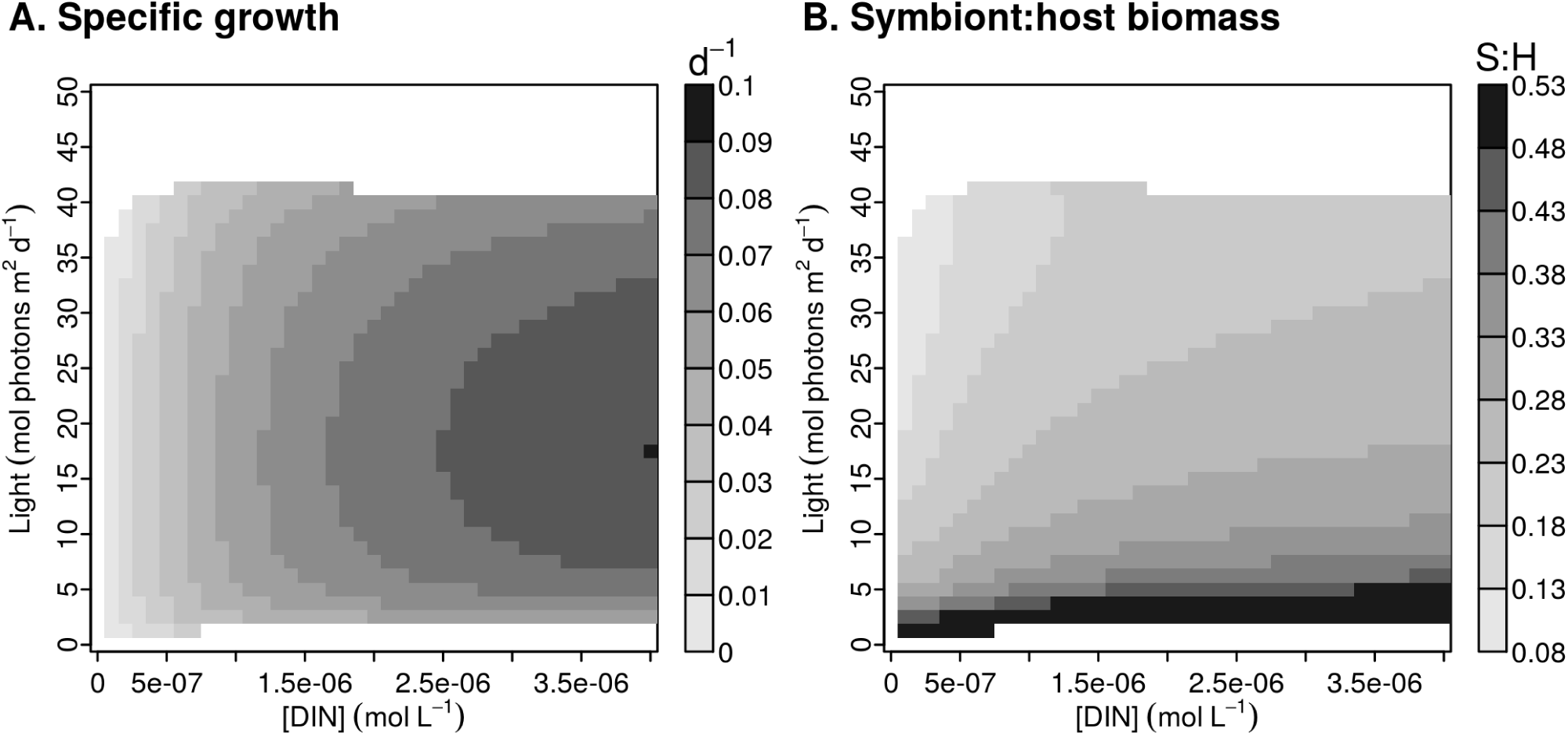
Steady state values of (A) specific growth (C-mol H C-mol H^-1^ d^-1^) and (B) the symbiont to host biomass ratio (C-mol S C-mol H^-1^) across gradients of external irradiance and dissolved inorganic nitrogen. Note that typical conditions for reefs are ∼1e-7 M DIN and 10-20 mol photons m^-2^ d^-1^. Simulations for each combination of light and nutrients (41 points along each axis) were run to steady state with all parameters at default values and prey density set to zero. Negative steady state growth rates and corresponding S:H ratios were set to zero, and a ceiling of 0.5 was imposed on S:H ratios to aid in visualization.

Similarly, increasing DIN alleviates nitrogen-limitation (Fig. 3A). Increased growth at higher DIN is predicted by the DEB model of Muller et al. (2009), and has also been observed experimentally (Muller-Parker et al. 1994; Tanaka et al. 2007; Tanaka et al. 2013). However, DIN elevation beyond a certain point (e.g., ∼3-4 μM in these simulations) has little effect on growth as carbon becomes limiting. Although very high nutrient levels may reduce growth in nature (Shantz, Lemoine, and Burkepile 2015), these impacts are not likely to occur within the range of concentrations considered here (<4 μM) (Ferrier-Pagès et al. 2000). In addition to increasing growth, DIN also increases the symbiont to host biomass ratio (Fig. 3B), a phenomenon also observed in reef corals (Marubini and Davies 1996). At low DIN and intermediate light, more typical of coral reef environments, symbiont to host biomass ratios are around ∼0.06-0.21, which is consistent with values reported in the literature (Muscatine, R McCloskey, and E Marian 1981; Edmunds et al. 2011; Hawkins et al. 2016).

The maximum predicted growth rates of ∼0.1 d^-1^, occurring between ∼10-25 mol photons m^-2^ s^-1^ light and ∼4 μM DIN (Fig. 3A), are comparable to the rate of 0.07 d^-1^ measured by Tanaka et al. (2007) in *Acropora pulchra* under similar N-enriched conditions. Under conditions more typical of reef environments (<0.5 μM DIN), predicted growth rates are ∼0.01-0.03 d^-1^. Observed specific growth rates in several coral species fall near or below the lower end of this range (∼0.01 d^-1^) (Osinga et al. 2011; Osinga et al. 2012), though values as high as 0.025 d^-1^ have been reported in *Galaxea fascicularis* (Schutter et al. 2010), and 0.04 d^-1^ in *Aiptasia diaphana*, a non-calcifying symbiotic anemone (Armoza-Zvuloni et al. 2014). However, it is not surprising that observed growth rates are often lower than model predictions, since the model does not account for ecological factors that may limit growth (e.g., competition, predation, bioerosion). Furthermore, while most measurements are made on skeletal growth, the model predicts biomass growth, which may not always be strongly correlated (Anthony 2002).

At irradiance levels above ∼25 mol photons m^-2^ d^-1^, steady state growth rates decline until positive growth ceases above ∼40 μmol photons m^-2^ d^-1^ (Fig. 3A). The mechanism underlying this decline is the increase in light energy beyond the capacities of photosynthesis and non-photochemical quenching: excess excitation energy generates reactive oxygen species (ROS) (Weis 2008; Roth 2014), which, in this model, have the phenomenological consequences of reducing the photosynthetic rate (representing photoinhibition) and increasing symbiont biomass loss (representing photodamage and/or symbiont expulsion) (see Eynaud, Nisbet, and Muller 2011). Together, these impacts reduce the symbiont to host biomass ratio (Fig. 3B), as occurs during coral bleaching. This reduction in symbionts consequently reduces the flux of fixed carbon to the host, resulting in increasing carbon-limitation (Fig. 3B) and eventual cessation of growth (Fig. 3A).

The incorporation of photooxidative stress in the model sets an upper limit to the amount of light at which a stable symbiotic interaction can be maintained, but even below this threshold of breakdown, negative effects of high light reduce steady state growth and symbiont:host biomass (Fig. 3). This gradual decline is consistent with experimental results showing that high light levels decrease growth (Schutter et al. 2011), and field studies documenting optimum growth rates at intermediate depths (Baker and Weber 1975; Huston 1985). By incorporating these impacts of light stress, the model predicts greater, and more realistic, variation in state variables across light gradients than was predicted by the models of Muller et al. (2009), which did not include photoinhibition or photodamage, or Eynaud, Nisbet, and Muller (2011), which included representations of photoinhibition or photodamage separately. It is important to recognize that the upper light limit set by photooxidative stress on a stable symbiosis under steady state conditions (Fig. 3) may be temporarily crossed by a dynamic system, which may experience a period of symbiont loss (bleaching) and reduced growth, after which a return to benign conditions may restore symbiont biomass and positive growth. To explore this further and illustrate the behavior of the model in more detail, we evaluate a number of dynamic simulations below (see “Dynamic behavior”).

## Sensitivity analysis

The values used for each parameter in the model (Table 2) are derived from relevant literature (see Supplementary Information). Here we evaluate the sensitivity of the model to changes in these parameter values, which also serves to demonstrate the behavior of the dynamical system. We calculated fractional change in steady state values in response to fractional changes in parameter values, relative to their default values, under environmental conditions typical of coral reefs.

Overall, relative changes in the steady state of the system are less than the equivalent relative change in parameter value. However, changes in certain parameter values have more significant impacts: increasing *j_Nm_* or decreasing *K*_*N*_ both dramatically increase host growth (Fig. 4), demonstrating the strong nitrogen-limitation that characterizes these symbioses. The parameter *ā*\* has a strong impact on S:H biomass ratios (Fig. 4) since this parameter determines the amount of light absorbed by symbionts, with lower values increasing light-limitation. Increasing the maximum growth and turnover rates have the expected effects of increasing and decreasing growth, respectively. Parameters relating to photooxidative stress and bleaching have little impact under low nutrients and intermediate light (Fig. 4), but have larger impacts under higher light (e.g., Fig. S6). Sensitivity analyses conducted under different combinations of external light and nutrients are presented in Figs. S3-S7.

**Figure 4:**
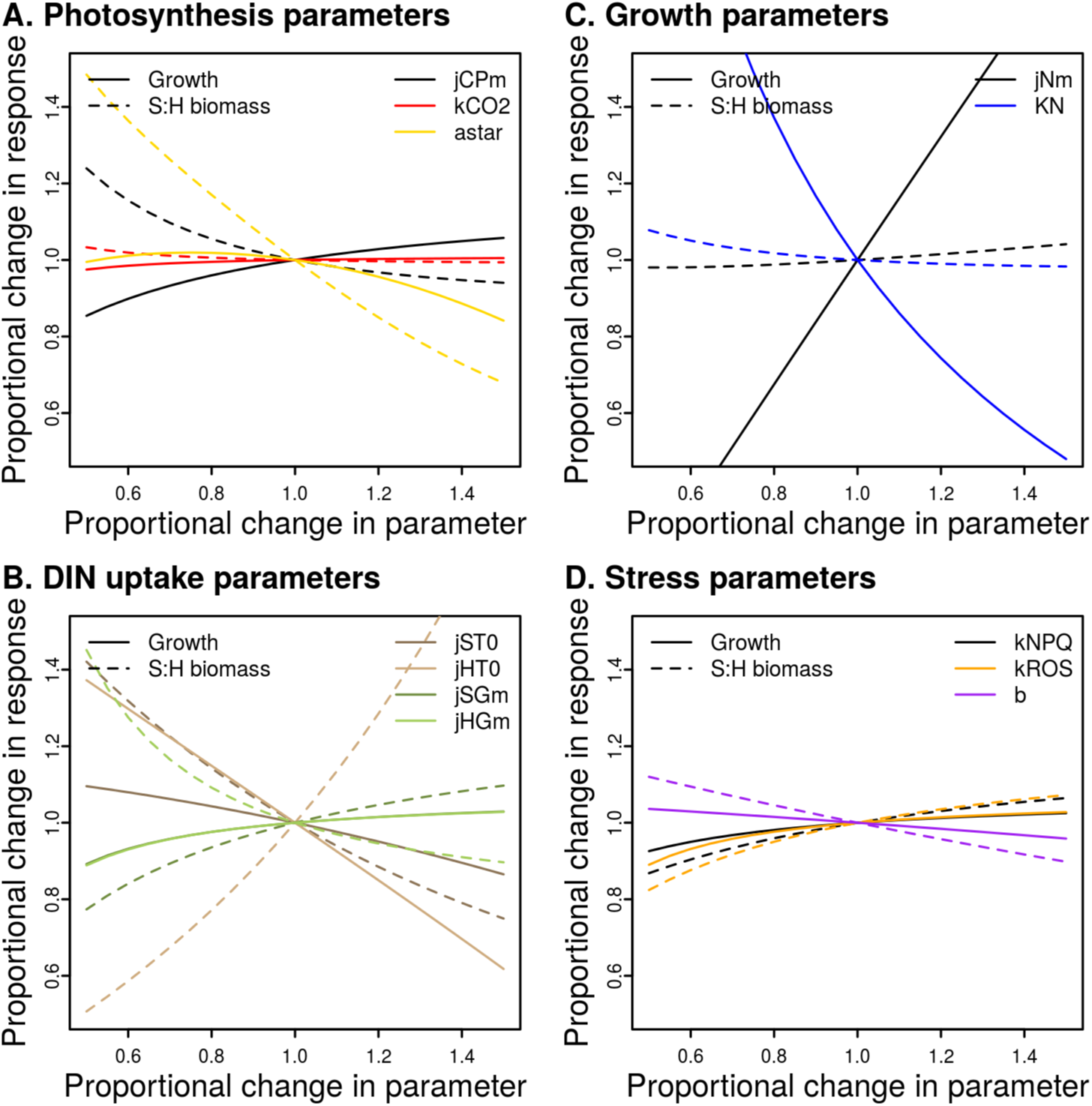
Sensitivity analysis. Plots show the fractional change in steady state values of growth (solid lines) and S:H biomass (dashed lines) in response to fractional changes in default parameter values (see Table 2 for default values). Parameters are grouped by the processes in which they are involved. This sensitivity analysis was conducted at conditions typical for coral reef environments: low DIN (1e-7 M) and intermediate light (15 mol photons m^-2^ d^-2^), with prey density set to zero. Sensitivity analyses conducted other environmental conditions are presented in Figs S3-S7.

## Dynamic behavior

The dynamic behavior of the model demonstrates its power to integrate multiple environmental forcings simultaneously. Here we present several scenarios that demonstrate the model’s ability to reproduce complex phenomena that have been observed in corals.

### Seasonal variability

Symbiont densities and coral growth rates are known to vary seasonally, representing an integrated response to changes in a suite of environmental factors. Light in particular is a strong driver of these trends (Stimson 1997; Brown et al. 1999; Fagoonee et al. 1999; Fitt et al. 2000), with high light associated with lower symbiont abundance and reduced tissue biomass. The role of light in driving seasonal changes in symbiont density was demonstrated nicely by Stimson (1997), who also found that experimental nutrient-enrichment amplified the light-driven seasonal oscillation. Using the levels of light and nutrients from this study as inputs, the model reproduces this observed interaction among environmental factors (Fig. 5), and also provides the mechanism: increasing light in summer decreases symbiont growth rates due to photooxidative stress, leading to decreasing S:H ratios. Under nutrient enrichment, this effect is more pronounced, as CO_2_-limitation of photosynthesis (due to higher symbiont standing stocks) causes mild bleaching that results in a similar summertime minimum S:H as the ambient nutrient case (‘physiological bleaching’ *sensu* Fitt et al. (2001)). Decreasing light into winter then alleviates the photooxidative stress constraints on carbon fixation such that nitrogen-limitation constrains the S:H ratio, explaining why S:H increases more when DIN is enriched (Fig. S8).

**Figure 5:**
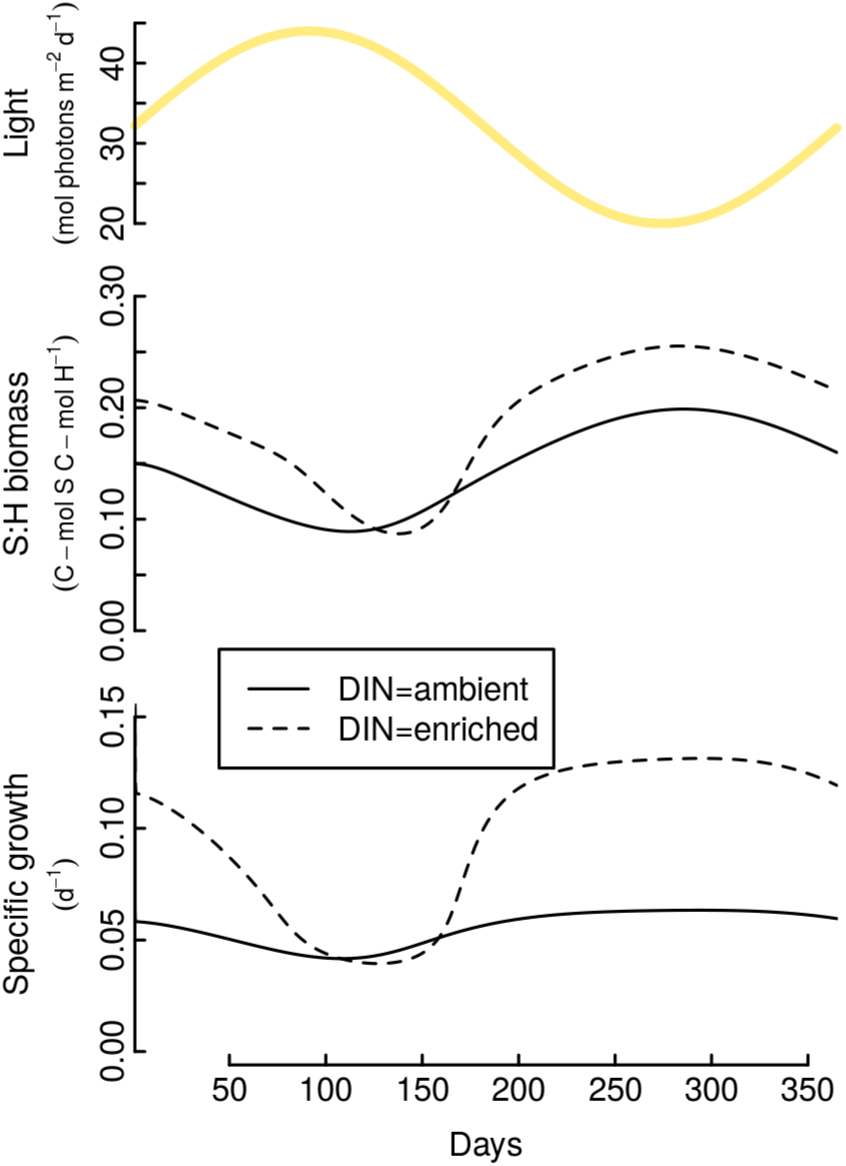
Light-driven seasonal dynamics of symbiont abundance and coral growth. Light input (upper panel) was designed as a sinusoidal curve with a period of one year, with maximum and minimum values of 44 and 20 mol photons m^-2^ d^-1^, corresponding to those measured by Stimson (1997). The dynamic behavior of symbiont to host biomass ratio (middle panel) and the specific growth rate of host biomass (lower panel) show seasonal oscillations that are greater in magnitude under high nutrients (15.14 *μ*M N; dashed lines) relative to low nutrients (0.14 *μ*M N; solid lines), consistent with the findings of Stimson (1997). Prey density was set at 1e-6 CmolX L^-1^.

The prediction of higher growth when light is reduced indicates that growth is not limited by low light in winter, but is actually reduced by excess light in summer^4^, consistent with the experimental findings of Schutter et al. (2011). Seasonal summertime reductions in tissue biomass have also been well-documented in the field (Fitt et al. 2000), along with reductions in net photosynthetic capacity (Muller-Parker 1987). Importantly, while light alone may drive seasonal dynamics in the ways discussed^5^, temperature fluctuations may attenuate or even reverse the effect of light as cooler winters depress metabolism; thus, the relative magnitude of fluctuation in temperature, light, and other factors may produce wide variability in the direction and magnitude of seasonal changes in growth and symbiont abundance, depending on location and microhabitat. Nevertheless, the seasonal variability predicted here (Fig. 5) is consistent with experimental and field observations for corals, and demonstrates the model’s ability to predict dynamic behavior that mechanistically integrates multiple environmental drivers.

### Coral bleaching and recovery

Coral bleaching is the stress-induced loss of symbiotic algae from coral tissues, which can occur in response to a variety of environmental stressors. In most cases, coral bleaching is thought to begin with photooxidative stress in symbiont photosynthesis (Lesser 1997), which triggers a cascade of events leading to symbiont expulsion (Weis 2008). As symbionts are expelled, the host receives less fixed carbon, which may then compromise its ability to activate CCMs that deliver CO_2_ to photosynthesis (Wooldridge 2013). Increasing CO_2_-limitation for remaining symbionts, along with an amplified internal light environment due to reduced self-shading (Enríquez, Méndez, and Iglesias-Prieto 2005), may further exacerbate photooxidative stress and accelerate symbiont expulsion, driving the coral into a bleached state.

While these positive feedbacks have been discussed previously in the literature, this is the first attempt to implement and explore their properties within a dynamical model. Interestingly, these feedbacks lead to alternate stable states in the symbiotic system. The ‘healthy’ state is characterized by nitrogen-limitation of both symbiont and host: under these conditions, the symbiont translocates sufficient carbon to support host growth and CCMs, which ensures that photosynthesis does not become CO_2_-limited. However, if carbon translocation is disrupted (and light is sufficiently high), then the system is driven into the ‘bleached’ state by photooxidative stress and positively reinforcing carbon- and CO_2_-limitation. In this context, coral bleaching can be understood as a transition from one stable state to another, and bleaching thresholds are sets of environmental conditions that push a healthy-state coral onto a trajectory leading to a bleached state.

We are highly interested in the conditions under which the system switches from a healthy to a bleached state, and can use this model as a tool to explore this dynamic behavior. Most straightforwardly, this switch occurs when increasing external irradiance (Fig. 6A) causes sufficient ROS production (Fig. 6B) and photoinhibition (Fig. 6C) that the positive feedbacks between host carbon-limitation (Fig. 6D) and CO_2_-limitation of photosynthesis (Fig. 6E) are rapidly engaged, leading to even greater photooxidative stress and a rapid decline in S:H biomass (Fig. 6F), characteristic of coral bleaching. However, the positive feedbacks involved in bleaching are not engaged only in response to high external irradiance alone; in fact, they depend on the relative balance of light energy absorption and quenching, which in turn depends on the availability of CO_2_ for photosynthesis. While previous models framed photooxidative stress as a fixed response to absolute external irradiance (Eynaud, Nisbet, and Muller 2011), our implementation considers the dynamic balance of multiple energy sinks in the causation of stress, which is more consistent with current understanding of symbiosis dysfunction (Wooldridge 2013), and establishes a critical role of host CCMs in providing CO_2_ for photosynthesis (Tansik, Fitt, and Hopkinson 2015; Hopkinson, Tansik, and Fitt 2015).

**Figure 6:**
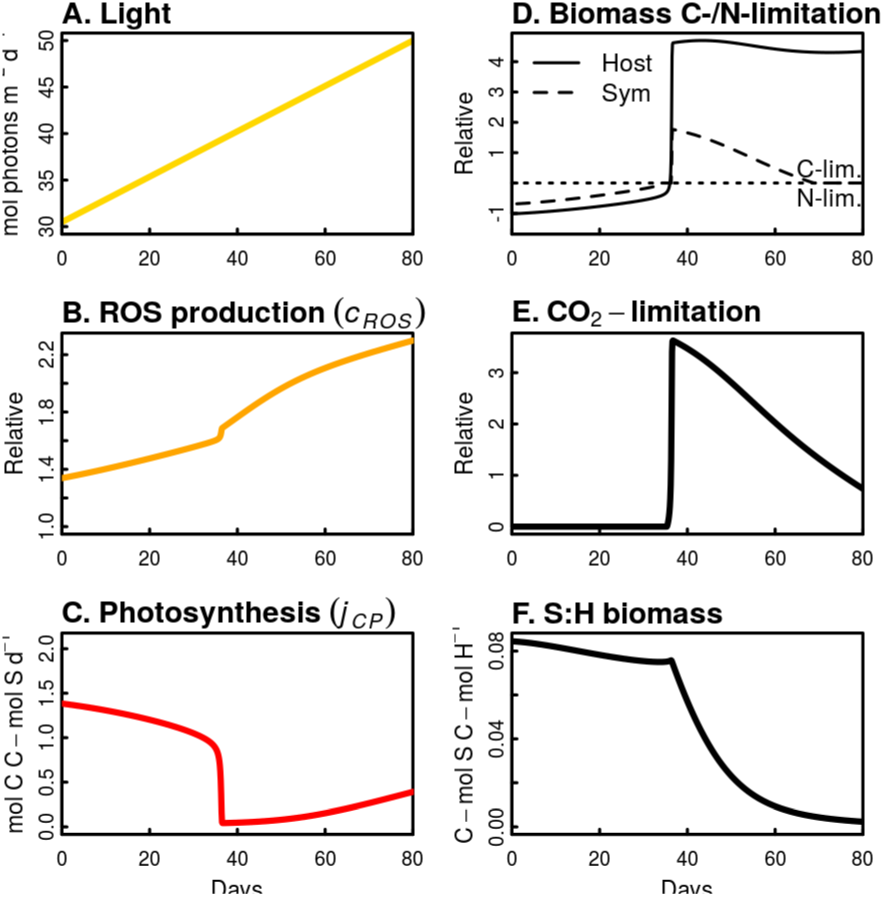
Coral bleaching as a switch from a nitrogen- to carbon-limited alternative stable state. This transition is demonstrated in response to gradually increasing external light (A), which causes an increase in production of ROS (B) that reduces the photosynthetic rate through photoinhibition (C). Decreasing photosynthesis moves the system from overall nitrogen-limitation toward carbon-limitation (D); when this threshold is crossed, the system rapidly becomes highly carbon-limited as photosynthesis becomes CO_2_-limited (E) and symbiont densities rapidly decline (F) into a bleached state. Substrate limitation coefficients were calculated using Eq. 23. All parameters were set at default values with external DIN=1e-7 mol N/L and prey density set to zero).

The importance of host CCM activity establishes significant interactive roles for other factors in influencing coral bleaching responses. For example, simulations of high light stress (Fig. 7) demonstrate that bleaching can be attenuated by heterotrophic feeding, a phenomenon which has been observed experimentally (Borell et al. 2008). The mechanism underlying this prediction is that feeding by the host increases host CCM activity, which delays the onset of CO_2_-limitation of photosynthesis and reduces bleaching severity. On the other hand, elevated nutrients exacerbate bleaching (Fig. 7), since higher symbiont densities are more susceptible to CO_2_-limitation (Wooldridge 2009). Several experimental (Wiedenmann et al. 2013; Cunning and Baker 2013; Vega Thurber et al. and correlational studies (Wooldridge and Done 2009) are consistent with this mechanistic link between high nutrients and bleaching.

**Figure 7:**
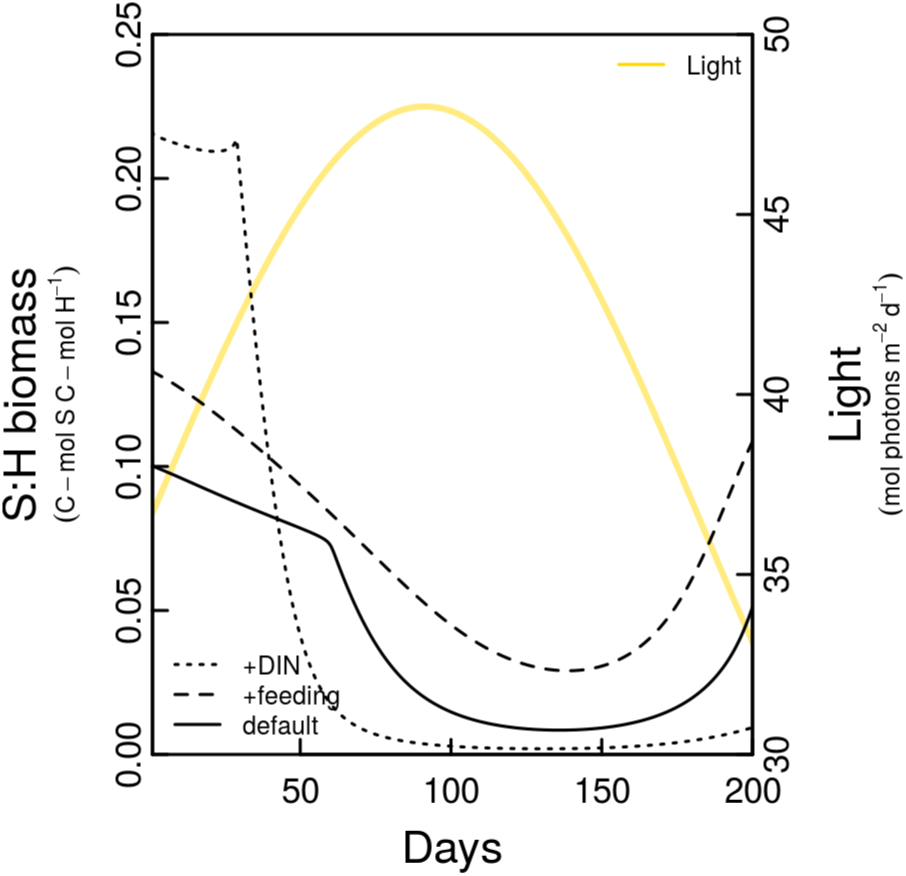
Bleaching with interactive factors. Simulations of high light stress (sinusoid with maximum=48 mol ph m^-2^ d^-1^) under default environmental conditions (solid line; DIN=1e-7 mol N L^-1^; prey=2e-7 C-molX L^-1^), or with elevated feeding (dashed line; prey=1e-6 C-molX L^-1^), or elevated nutrients (dotted line; DIN=4e-6 mol N L^-1^). Initial symbiont biomass was set to the steady state for each set of starting conditions, with all other parameters at default values.

Since bleaching can be understood as a transition from a nitrogen-limited state with high S:H biomass to a carbon-limited state with low S:H biomass, induced by an external stressor, recovery can be understood as a switch back to the nitrogen-limited state once the external stressor is alleviated. In natural settings, this typically occurs through seasonal declines in temperature and light. However, hysteresis associated with the system’s alternate stable states indicates that the symbiosis cannot recover along the same trajectory it followed during bleaching; indeed, the stressor must be alleviated *below* the threshold that initially caused bleaching in order for the system to recover (Fig. 2). This is because under the same external conditions, a bleached coral with low S:H biomass (relative to a healthy coral with high S:H biomass) is characterized by greater light amplification and weaker CCM activity, which enhance photooxidative stress and serve to maintain the carbon-limited state. In order for the system to recover, the stressor must be reduced enough such that photooxidative stress ceases and translocation of carbon from symbiont to host is resumed. Once this occurs, the host can energize its CCMs, which further enhances carbon fixation and accelerates the system back toward a nitrogen-limited state, indicative of recovery. The conditions under which recovery can occur – which determine the magnitude of hysteresis (Fig. S9) – depend on the relative abundance of nitrogen and carbon in the environment. Higher food levels, representing a non-autotrophic carbon source for the host, make it easier for the host to overcome carbon-limitation, thus providing a potential mechanism underlying observations that feeding aids recovery from bleaching (Grottoli, Rodrigues, and Palardy 2006; Connolly, Lopez-Yglesias, and Anthony 2012). Conversely, high external DIN impedes the re-establishment of nitrogen-limitation, making recovery from bleaching more difficult (i.e., narrowing the sets of conditions under which recovery is possible).

Dynamic simulations of recovery reveal another interesting behavior of the system: under some conditions, an ‘overshoot’ occurs in which S:H biomass temporarily increases beyond the ratio maintained in the ‘healthy’ state, before returning to stabilize at this value (Fig. 8). In fact, unusually high symbiont densities have been observed following bleaching in both experimental (Cunning, Silverstein, and Baker 2015) and field studies (Kemp et al. 2014), and have been interpreted as a potential ‘disequilibrium in host-symbiont regulation’ (Kemp et al. 2014). The model reveals the dynamics of this ‘overshoot’ as follows: as symbionts repopulate the host, photosynthesis becomes increasingly CO_2_-limited due to weak CCM activity of the carbon-limited host. Thus, although symbiont growth is not yet carbon-limited, a growing symbiont population has less and less excess carbon (per symbiont) to share, and is thus moving toward carbon-limitation. Meanwhile, because S:H biomass is increasing, the host receives more and more carbon per unit host biomass, and is thus moving away from carbon-limitation. If the host overcomes carbon-limitation *before* the symbiont reaches it, then the system rapidly transitions to the nitrogen-limited state and the S:H ratio stabilizes without an overshoot (Fig. 8A). However, if the symbiont becomes carbon-limited first (Fig. 8B, 8C), then carbon translocation per symbiont declines further and photosynthesis becomes more CO_2_-limited, which maintains carbon-limitation of the host. In this situation, continued growth of less and less productive symbionts drives S:H biomass to a much higher level before the host finally overcomes carbon-limitation. At this point, representing the peak of the overshoot, the transition to nitrogen-limitation finally takes place and the S:H biomass ratio declines and stabilizes as positive growth is resumed.

**Figure 8:**
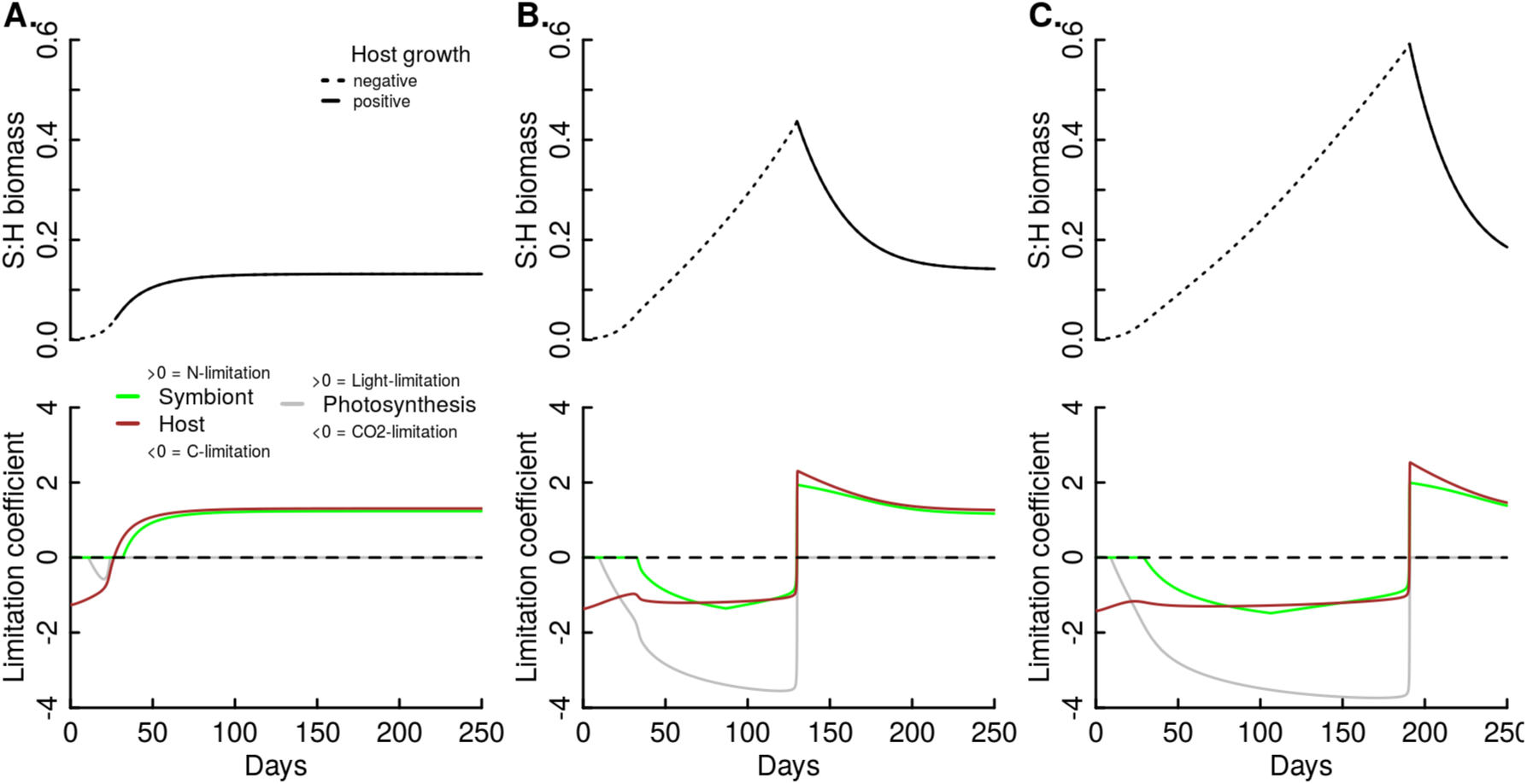
Recovery from bleaching with varying N:C availability (A: N:C=0.100; B: N:C=0.175; C: N:C=0.220). Higher N:C ratios in a heterotrophic food source (effectively representing higher external DIN and/or lower feeding rates) cause a larger overshoot in the S:H biomass ratio, and prolong the duration of time until the system ‘recovers’ by re-establishing nitrogen-limitation. Substrate limitation coefficients were calculated using Eq. 23. All simulations were run with default parameters (except varying *n*_*NX*_), with L=20 mol photons m^2^ d^-1^, DIN=1e-7 mol N L^-1^, prey=1e-7 CmolX L^-1^, and initial S:H biomass=0.001.

Whether this ‘overshoot’ occurs or not is determined by the relative availability of carbon and nitrogen to the host – any factor that enhances carbon-limitation of host growth (e.g. high DIN and/or low feeding) therefore magnifies the overshoot and prolongs the dysfunctional, carbon-limited state of the symbiosis (Fig. 8). On the other hand, factors that favor nitrogen-limitation, such as low external DIN and/or high feeding rates, will accelerate the re-establishment of nitrogen-limitation and prevent an overshoot from occurring at all. While many scenarios are possible under different environmental conditions, we illustrate the general effects of varying N and C availability on recovery from bleaching with a series of simulations that vary the N:C ratio of host’s heterotrophic food source (Fig. 8A-C): lower N:C ratios (effectively representing lower DIN and/or higher heterotrophy) favor nitrogen-limitation and more rapid recovery, while higher N:C ratios (effectively representing higher DIN and/or lower heterotrophy) favor carbon-limitation and prolonged recovery with a larger overshoot in S:H biomass.

These dynamics reveal that the re-establishment of nitrogen-limitation is the most important diagnostic of recovery to a ‘healthy’ state, as this is when positive growth rates are resumed. A high S:H biomass ratio alone does not necessarily indicate that a ‘healthy’ state has been reached, since the carbon-limited state may still persist (e.g. Fig. 8B, 8C). This could explain why corals that have recovered their symbiont populations after bleaching may still exhibit energetic deficits and physiological impacts, possibly for months to years (Levitan et al. 2014, Hughes and Grottoli (2013))). These findings suggest that host acquisition of carbon from a source other than the symbiont may be extremely important for the system to recover from bleaching. Indeed, host feeding has been shown to promote a more rapid return to pre-bleaching levels of key physiological parameters in recovering corals (Connolly, Lopez-Yglesias, and Anthony 2012). Additional carbon sources for the host, such as direct uptake of dissolved organic carbon (Levas et al. 2015), may also promote more rapid recovery from a bleached state.

## Conclusions

This dynamic bioenergetic model of coral*-Symbiodinium* symbioses mechanistically reproduces patterns in steady-state coral growth and symbiont abundance commonly observed in corals, including higher symbiont abundance with higher nutrients and feeding, lower symbiont abundance with increasing light, and optimal growth at intermediate light levels. Moreover, the model reproduces complex dynamic behaviors including seasonal changes in symbiont density at different nutrient levels, rapid bleaching above a threshold of high light, mitigation of bleaching by heterotrophic feeding, exacerbation of bleaching by elevated nutrients, and an overshoot of symbiont density during recovery from bleaching. These examples demonstrate the model’s ability to integrate multiple environmental forcing functions to reproduce complex responses; meanwhile, the diversity of these phenomena suggest the model has captured many of the important features of the system in a unifying mechanistic framework. This model also provides a new conceptual framework for considering coral bleaching as a transition to an alternate stable state, which has important implications for understanding the performance and maintenance of symbiotic interactions. In this context, the ‘healthy’ stable state represents a scenario in which nitrogen-limitation stabilizes the symbiont to host biomass ratio and maintains positive growth. Conversely, carbon-limitation represents a dysfunctional state wherein positive feedbacks result in a loss of photosynthetic capacity and negative growth. Importantly, these ‘control’ mechanisms arise entirely passively through feedbacks present in the system.

In addition to the suite of specific examples presented here, the model can be used to explore many different dynamic environmental scenarios, and represents a tool that biologists and ecologists can use to generate hypotheses and make predictions in both experimental and natural settings. Moreover, parameter values can be modified to correspond to different genetic or functional types of coral hosts and *Symbiodinium* partners in order to evaluate variability in system responses. Thus, the diversity of potential applications for this model is high, and we envision this work as a foundation for continued development, which may include more detailed treatments of specific modules (e.g., DIC processing), and the incorporation of more external forcing capabilities (e.g., external DIC, temperature). Importantly, open source R code allows this effort to benefit from contributions from the wider scientific community, including those with empirical and theoretical backgrounds. Ultimately, the continued refinement of these tools is fundamental in elucidating the mechanisms of symbiosis function and dysfunction, and in predicting coral responses to environmental change.

## Acknowledgements

This work was supported by a NSF Postdoctoral Research Fellowship in Biology to RC (#1400787).

## Footnotes

1 Rejection fluxes must always be positive, and hence are specified with the notation (*x*)_+_, which means *max*(*x*,0).

2 Note that recycling of symbiont biomass turnover (*rNS* and *rCS*) only occurs based on the maintenance component of turnover (i.e., 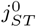), and not the photodamage/bleaching component, as this loss represents biomass that is expelled from the host.

3 See footnote 2.

4 at the light levels indicated in Stimson (1997). Note that if light levels were reduced throughout the year (e.g. for a coral at greater depth) such that light did not cause photo-oxidative stress in summer but became limiting to photosynthesis in winter, the S:H ratio would still increase in winter, but growth would decrease; the latter scenario is predicted both by the present model as well as that of Muller et al. (2009), since photo-oxidative stress does not become relevant.

5 See footnote 4.

